# Phenotypic cliffs in the RNA genotype-phenotype map

**DOI:** 10.1101/2025.10.06.680761

**Authors:** Paula García-Galindo, Sebastian E. Ahnert

## Abstract

Point mutations of a genotype can leave the phenotype unchanged, or change it, in some cases radically. The extent of this phenotypic change can critically impact fitness. To investigate the range of possible phenotypic changes that result from point mutations, we analyse the structure of the well-established RNA genotype–phenotype (GP) map for sequences of length 12 by developing a general phenotypic distance framework. Our analysis reveals that phenotypes are more likely to be surrounded by similar phenotypes than would be expected by chance. Furthermore, we see that generalised versions of the GP map metrics of frequency, robustness, and evolvability that take phenotypic distance into account still exhibit the same fundamental structural relationships that are observed in many GP maps. We also develop site-specific quantities for robustness, evolvability, and accessible phenotypic distance, which reveal a key insight: RNA secondary structure (SS) sites that are more likely to be neutral and access fewer new phenotypes tend to produce larger changes in phenotype when a change does occur. Robust sites therefore produce cliffs in the landscape (flat in some directions, steep in others) whereas non-robust sites give rise to more gently undulating landscapes.

## I. INTRODUCTION

The quantitative study of phenotypic variation at the molecular level can illuminate differences in biological function and evolutionary histories [1]. We employ comprehensive RNA sequence-to-structure mapping to quantify structural differences between folded RNAs in order to shed light on the range of phenotypic changes that mutations can produce.

The sequence-to-structure map of short RNA is computationally tractable and biologically relevant, as the biological functions of short RNA (e.g., tRNAs, miRNAs) are often dependent on their folded structure [2, 3]. This map takes the RNA nucleotide sequence as the genotype, which is subject to mutations, and the resulting RNA fold as the phenotype, which is subject to natural selection. For short RNAs, the three-dimensional tertiary structure is wellapproximated by the secondary structure fold [4]. As a result, the GP map of RNA of length *L* represents all 4^*L*^ possible *L*-sequences mapped to their respective secondary structures. Typically the genotype space is represented as a network in which the nodes are individual sequences, and the edges correspond to the possible point-mutations between them. [4].

RNA is not a static biopolymer; due to thermal fluctuations at the molecular scale, an RNA sequence can stochastically shift between different folds [5, 6]. Instead of only considering one phenotype per genotype like in the typical deterministic GP map, which maps the minimum free energy structure, the structural plasticity of RNA can be accounted for by defining a many-to-many GP map known as the non-deterministic genotype-phenotype (ND GP) map, which assigns the Boltzmann ensemble of probable secondary structures as the phenotype [7]. In the following sections we will mainly consider phenotypic distance in deterministic GP maps, but also show that it can be readily generalised to ND GP maps.

We define phenotypic distance in the context of RNA secondary structures by applying the Hamming distance to the secondary structure (SS) dot-bracket notation [8]. In this notation, base pairs are denoted by matching parentheses, and unpaired bases are represented by dots. For example, the sequence ACGCCUCGGGGA corresponds to the secondary structure .(.(….).)., which contains two Watson-Crick GC base pairs [4]. To compare two RNA SS of the same length *L*, we use the Hamming distance *H*(*p, q*) and the similarity *S*(*p, q*):

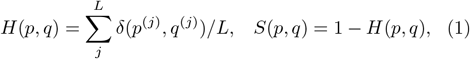

where *p*^(*j*)^ and *q*^(*j*)^ represent the symbols at position *j* in the dot-bracket notations of structures *p* and *q*, respectively.

The Hamming distance allows us to assess how mutations impact phenotypic variation by comparing the secondary structure at each sequence position in the dot-bracket notation. The phenotypes that differ more from the wild type are those with a higher Hamming distance *H*(*p, q*) or lower similarity *S*(*p, q*).

We incorporate phenotypic distance into the structural analysis of GP maps by developing a framework of key GP map quantities that is weighted by the Hamming distance. These quantities are: (a) frequency, which is the number of genotypes mapping to a certain phenotype; (b) robustness, how insensitive a phenotype is to mutation; and (c) evolvability, the number of novel phenotypes that are discovered through mutation. We apply the framework to the deterministic GP map of RNA secondary structures of length *L* = 12, and the corresponding ND GP map. We observe that several universal structural properties [4, 9] of the GP map are preserved. Specifically these are *bias*, which describes a highly skewed distribution in the number of genotypes per phenotype; *neutral correlations*, which means that genotypes of the same phenotype are separated by fewer mutations than expected by chance; and *correlated robustness and evolvability*, which relates to the negative correlation of genotypic robustness and evolvability and the positive correlation of phenotypic robustness and evolvability.[4, 9]

This framework also allows us to systematically analyse distributions of phenotypic distance, both in the local context of point-mutation neighbourhoods and the global context of the entire GP map. We also analyse the phenotypic distance distribution for mutations at a specific site to gain a deeper understanding of the effect of mutation on phenotypic change, and the relationship between phenotypic distance and the more established metrics of and evolvability and robustness. We find, on average, a positive correlation between site-specific robustness and average sitespecific Hamming distance. This suggests that mutations at robust sites tend to either preserve the phenotype or, when they do result in change, lead to phenotypes that are substantially more different than those arising from mutations at less robust sites. We suggest that this adjacency of ‘flat’ and ‘steep’ directions of phenotypic change in the mutation landscape can be visualised as a ‘phenotypic cliff’.

## II. METHODS

### A. Weighted phenotypic frequency

The phenotypic frequency of a phenotype *p* is the number of genotypes that map to that phenotype, *F*_*p*_, divided by the total number of possible genotypes, *f*_*p*_ = *F*_*p*_*/K*^*L*^. It represents the probability that we pick structure *p* if we randomly sample the genotype space [4].

We define the *weighted phenotypic frequency* of *p*, 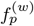, as the frequency of phenotypes weighted by their similarity to *p*, in other words, it represents the global average similarity to *p* across the entire genotype space:

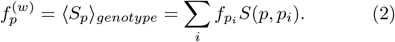

Instead of only accounting for the genotypes that map to *p*, the weighted phenotypic frequency therefore considers all genotypes and weights their contributions according to the similarity of their phenotypes to *p*. By definition the weighted phenotypic frequency will therefore exceed the conventional, unweighted phenotypic frequency, 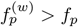, and the additional contributions will highlight the degree to which structures similar to *p* are present across the GP map.

### B. Weighted generalisations of robustness and evolvability

Two key quantities often used to systematically analyse GP map structure are *evolvability*, which quantifies the potential for phenotypic change, and *robustness*, which measures the degree to which mutations leave a phenotype unchanged [10–12]. These metrics are adapted to the phenotypic distance framework by incorporating the similarity *S* and Hamming distance *H*, thus introducing weighted quantities based on structural difference between the reference phenotype *p* and other phenotypes *p*^*′*^ in the GP map.

The genotypic robustness *ρ*_*g*_ of a genotype *g* with phenotype *p* represents the proportion of 1-mutation neighbors of *g* that also have phenotype *p* [12]. In the weighted frame work this becomes 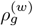, which corresponds to the average similarity of phenotypes to *p* across the 1-mutation neighbours of *g*, denoted as 𝒩_*g*_:

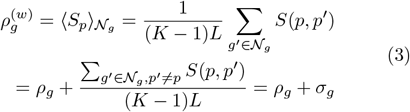

where the term summarised as *σ*_*g*_ only accounts for the similarity of non-neutral phenotypes *p*^*′*^ ≠ *p* in the neighbourhood of *g*, so that we can separate this contribution from the unweighted robustness *ρ*_*g*_.

The phenotypic robustness *ρ*_*p*_ is defined as the average of *ρ*_*g*_ over all sequences that map to *p* in the unweighted case [12]. In the weighted framework this becomes the average similarity to *p* over 𝒩_*p*_, the 1-mutation neighbours of all genotypes that map to *p*:

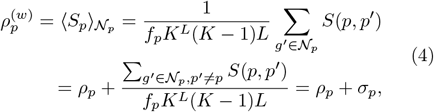

The *σ*_*p*_ expression again only accounts for the similarity of non-neutral phenotypes *p*^*′* ≠^ *p* in the neighbourhood of *p*, so that we can separate the unweighted robustness *ρ*_*p*_ from the weighted contribution. In GP maps he concept of evolvability has been defined as the number of uniquely different phenotypes accessible in the 1-mutation neighbourhood of a genotype or phenotype [7, 12], which we will refer to as “discovered phenotypes”. Each unique structure is only counted once even if it maps to multiple sequences in the mutational neighbourhood under consideration. We can incorporate phenotypic distance into a weighted generalisation of evolvability by considering the set of all uniquely different phenotypes, and summing the fractions of their differences to *p*, which corresponds to the sum over the Hamming distances between these phenotypes and *p*.

Let us denote as 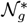 the set of *unique* phenotypes 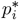 that map to the genotypes in the 1-mutation neighbourhood 𝒩_*g*_ of a sequence *g* with structure *p*. The unweighted genotypic evolvability is then 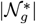. The weighted genotypic evolvability is therefore the sum of the Hamming distances between *p* and the 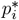 in 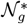

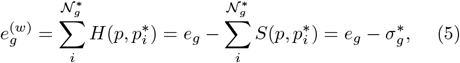

where the 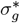 represents the sum over the similarities of the discovered phenotypes in the neighbourhood of *g*. In the same way, at the phenotypic level, the phenotypic evolvability sums the Hamming distances between *p* and all unique structures in 𝒩_*p*_, the 1-mutation neighbourhood of all genotypes that map to *p*:

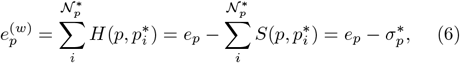

where the 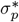 again represents the sum over the similarity of the discovered phenotypes in the neighbourhood of *p*. In this framework, a large number of uniquely different structures does not by itself guarantee a high evolvability, as the value of the evolvability also depends on the phenotypic difference.

As the *σ* contributions in Eq.3–6 are all positive the phenotypic distance framework defines a weighted robustness that is greater than the unweighted one and a weighted evolvability that is smaller than the unweighted one.

### C. Generalisations to the non-deterministic (ND) GP map

Recent work redefines frequency, robustness and evolvability for a non-deterministic (ND) GP map, in which each genotype maps to a probability distribution of structures. Such ND GP maps are arguably more realistic, as RNA sequences can fold into multiple different structures [7]. These maps can also be used to study RNA plasticity [13]. In the non-deterministic map quantities such as frequency, robustness, and evolvability are defined as probabilistic ensemble averages [7], such that each nondeterministic quantity 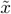 corresponds to an average over all the possible ND GP map realisations of the *x* deterministic quantity: 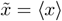. We can incorporate phenotypic distance into ND GP maps the same manner as in the deterministic case, where the quantities are weighted by similarity or Hamming distance. However, the non-deterministic case measures the presence of a phenotype wi ity mass. For example, the weighted non-d quency of *p* is the average similarity to *p* o probability mass in genotype space:

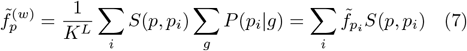

We follow the same definition for the deterministic robustness and weighted nondeterministic robustness and weighted non-deterministic evolvability. To validate each definition n we compute the ensemble average and check its correspondence to the weighted non-deterministic quantity, approaching 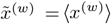, which is shown in the Supplement for the case of the RNA12 GP map.

## III. RESULTS

### A. The weighted RNA12 GP map res GP maps in terms of key structural properties

In many GP maps a negative correlation of genotypic robustness and evolvability can be observed [4, 12] because there is a fixed number of mutational neighbours for any genotype. If more mutants are robust, fewer are able access new phenotypes. This relationship is also present in the weighted RNA12 framework, where we observe a negative correlation between the weighted genotype robustness (Eq.3) and weighted genotypic evolvability (Eq.5), shown in Fig.1A with Pearson *r* : −0.59.

**FIG.1.**
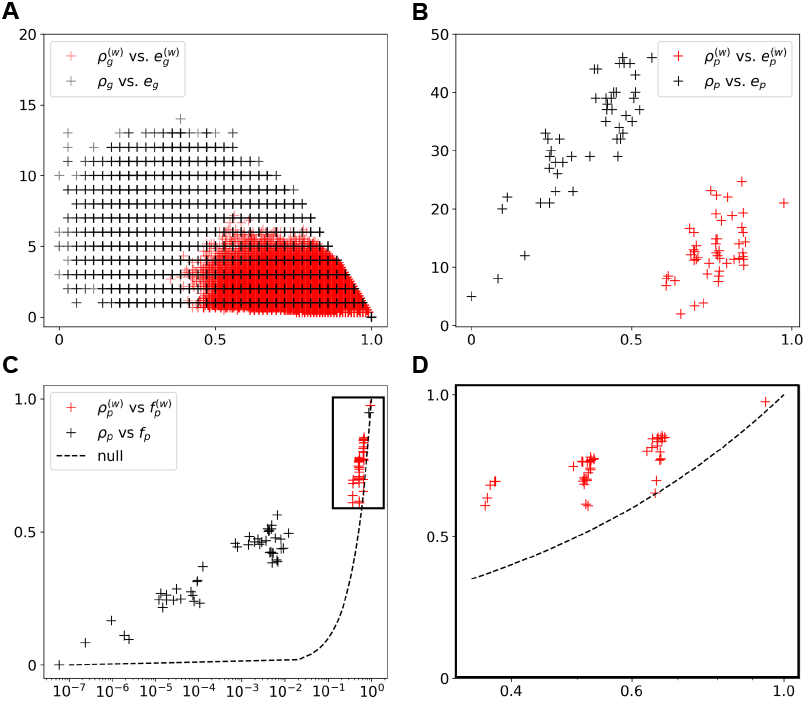
Structural properties are present in the phenotypic distance framework. **(A)** Negative correlation between the genotypic robustness and evolvability for the weighted (Pearson *r* : 0. *™* 59) and non-weighted case.**(B)** Positive correlation between the phenotypic robustness and evolvability for the weighted (Pearson *r* : 0.49) and non-weighted case. **(C)** The presence of genetic correlations is verified through the scaling *ρ*_*p*_ ~ log*f*_*p*_ for both the non-weighted and **(D**, enlarged excerpt from **C)** weighted case.

At the phenotypic level, a positive correlation between phenotypic robustness and evolvability has been demonstrated across many GP maps [4, 12]. This in part because larger neutral spaces are more robust, and also adjacent to a larger number of alternative phenotypes[9, 12], but also hinges on non-local effects of mutations on sequence constraint [4, 14, 15], meaning that mutations in one sequence position can affect the degree to which sequence elsewhere is constrained or unconstrained. In an RNA context this would apply to the mutation of one base in a Watson-Crick pair, which would change the constraint of the paired base elsewhere in the sequence. In a genetic context one might consider the effect of mutating a stop codon and thereby changing the reading frame and thus the resulting change of a sequence from coding to non-coding or vice-versa. We also find a positive correlation of phenotypic evolvability and robustness in the weighted RNA12 framework as shown in Fig.1B of Pearson *r* : 0.49, thus replicating this widely observed relationship.

Neutral correlations are present in a GP map if genotypes that map to the same phenotype are more likely to be mutationally adjacent than one would expect based on their frequency. [9]. It has been previously shown that for RNA secondary structure and other GP maps these neutral correlations are strong, the robustness follows the scaling relationship *ρ*_*p*_ ~ log*f*_*p*_[4, 9, 14]. In the case of weighted phenotypic robustness and frequency, the relationship is still present 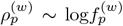 (Fig.1C,D). This means that, like in the unweighted case, similar phenotypes are on average closer in genotype space compared to the null model which in the weighted case would define the average similarity over the neutral space as being equal to the average similarity over the full genotypic space: 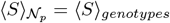.

### B. Phenotypes are surrounded by more similar phenotypes than expected by chance

We next examine the distribution of phenotypic changes that a single mutation can cause, by defining a weighted phenotype mutational probability.

The standard, unweighted phenotype mutational probability *ϕ*_*pq*_ is defined as the likelihood that a phenotype *q* mutates to phenotype *p*. This is a generalisation of robustness and is computed by taking the proportion of 1-mutation neighbors of phenotype *q* that lead to *p* [9, 16]:

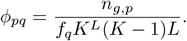

To define a phenotype mutation probability that incorporates phenotypic distances, we use the Hamming distance and define the weighted phenotype mutation probability *ϕ*_*Hq*_, which is the likelihood that a phenotype *q* mutates to a phenotype of Hamming distance *H*:

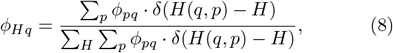

which is the general formula that simplifies to:

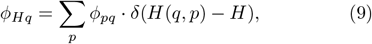

if we include all of the phenotypes *p* that *q* could possibly access, including the deleterious phenotype (for RNA this is the unfolded structure [4]), since ∑_*p*_ *ϕ*_*pq*_ = 1. We can also define the average weighted phenotype mutation probability for a given Hamming distance as *ϕ*(*H*) = ⟨ *ϕ*_*Hq*_ ⟩_*q*_. To define the global weighted phenotype probability Φ_*Hq*_, which is equivalent to the frequency of phenotypes a distance *H* from *q*, the probability is calculated over the genotype space (null model [9]) instead of the 1-mutation neighbours of phenotype *q*:

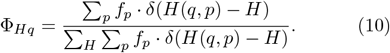

Here too we can define a global quantity Φ(*H*) = ⟨Φ_*Hq*_⟩_*q*_. The average phenotypic distance mutation probabilities are shown in Fig.2 for the RNA12 GP map. The probabilities to all 1-mutational neighbours, including the deleterious phenotype (Eq.9) are shown in Fig.2A. The probabilities including all 1-mutations except to and from the deleterious are shown in Fig.2B. Finally, Fig.2C just includes the 1-mutations to and from phenotypes that are non-deleterious and that change the phenotype, so we exclude the robustness term, when *H* = 0. In all cases we observe a bias towards robustness (*H* = 0) and similar structures (low *H >* 0) in the local neighbourhood (black), as given by *ϕ*(*H*), compared to the global distance distribution (blue) given by Φ(*H*), such that the GP map structure connects phenotypes that are similar to each other than expected by chance, from random sampling of a phenotype from the genotype space (Eq.10).

**FIG.2.**
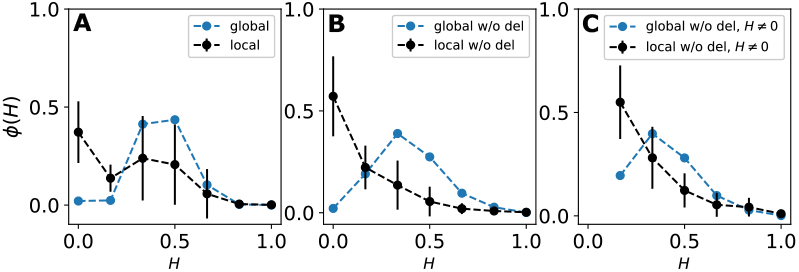
Phenotypes are on average more likely to be surrounded by similar phenotypes than expected by chance. Here we show the average probability *ϕ*(*H*) that a phenotype in the RNA12 GP map mutates to another phenotype that is different by Hamming distance *H* (black) as well as the probability Φ(*H*) of finding a phenotype of distance *H* if we sample from the full genotype space. In **(A)** the probability function considers all possible 1-mutation neighbours and follows Eq.9. In **(B)** The probability function considers all possible 1-mutation neighbours except to and from the deleterious phenotype (unfolded structure for RNA [4]), and follows Eq.8. **(C)** The probability function considers all possible 1-mutation neighbours except to and from the deleterious phenotype (unfolded structure for RNA [4]) and the robustness term, meaning those that do not change the phenotype, *H* = 0.

This verifies the phenotypic similarity correlations that were previously found. These results are expected from previous studies on RNA secondary structure ensembles, which show that mutational robustness of an MFE structure correlates with its structural similarity to other structures within the phenotype ensemble [17], and how an MFE structure correlates to the suboptimal structures of its 1-mutation neighbours, a phenomenon called plastogenetic congruence [6], and how this can be translated to predictions of phenotypic changes for random mutations [18]. This relationship shows that high phenotypic distance phenotypes *H* ≥ 0.5 are unlikely to be accessed through 1-mutations. We next investigate how we can access these unlikely and large phenotypic changes.

### C. Mutations of robust sites result in large phenotype changes, and thus in ‘phenotypic cliffs’

Computational studies of a range of different GP maps have consistently revealed similar structural properties of these maps [4], which we also verified for the weighted RNA12 GP map in Section III A. A possible explanation for the seeming universality of these properties can be found in the commonly observed organisation of genotypes into evolutionarily constrained and unconstrained parts [14]. Constrained sites are more sensitive to mutations and will lead to a more immediate change in phenotype, while unconstrained sites can mutate more freely without directly affecting the phenotype. For example, in RNA the paired nucleotides (represented by brackets in the dot-bracket notation) are more constrained relative to the unpaired nucleotides (represented by dots) [19]. We compare the degree to which mutations of unconstrained sites give rise to phenotypic change compared to mutations of constrained ones, by first defining site-specific structural quantities and phenotype mutation probabilities. The site-specific phenotypic robustness measures the number of mutations leading back to *p* when mutating genotypes at a specific site *i*:

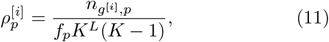

and the site-specific phenotypic evolvability counts the number of uniquely different accessible phenotypes that can only be accessed from site *i*:

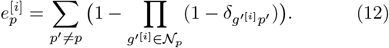

To check the likelihood of accessing a phenotype *p* from a particular site we can compute the site-specific phenotype mutation probability:

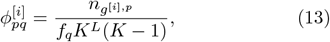

which can be generalised to a weighted site-specific phenotype mutation probability, 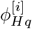, in the same way as for Eq.8. Finally, the site-specific average Hamming distance that a phenotypic site can access is:

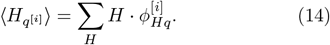

To see the distributions of phenotypic changes that arise from a mutation at a given site, we compute the site-specific phenotypic robustness, site-specific fractional evolvability, and site-specific average Hamming distance 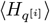 for each secondary structure phenotype on the RNA12 GP map and examine their relationship. We illustrate these quantities for a high frequency phenotype in Fig.3A. The unconstrained sites, which have high site-specific phenotypic robustness 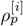 exhibit larger phenotypic changes (larger 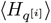, shown in black) than the constrained, while the sitespecific evolvability (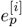, shown in red) is lower for these unconstrained sites than for the constrained. In Fig.3B we show the distribution of Hamming distances that are accessible from two different sites, site 5 which is unconstrained, and site 10 which is constrained. The weighted phenotype mutation probability distribution in Fig.2C resembles the distribution of Hamming distances for the constrained site Fig.3B (bottom), suggesting that constrained sites dominate in the average weighted phenotype mutation probability distribution for RNA12, and thus mask the somewhat different distributions that occur through mutations of unconstrained sites, which are wider and are centred around larger phenotypic distances of *H ≥* 0.5 as shown in Fig.3B (top).

**FIG.3.**
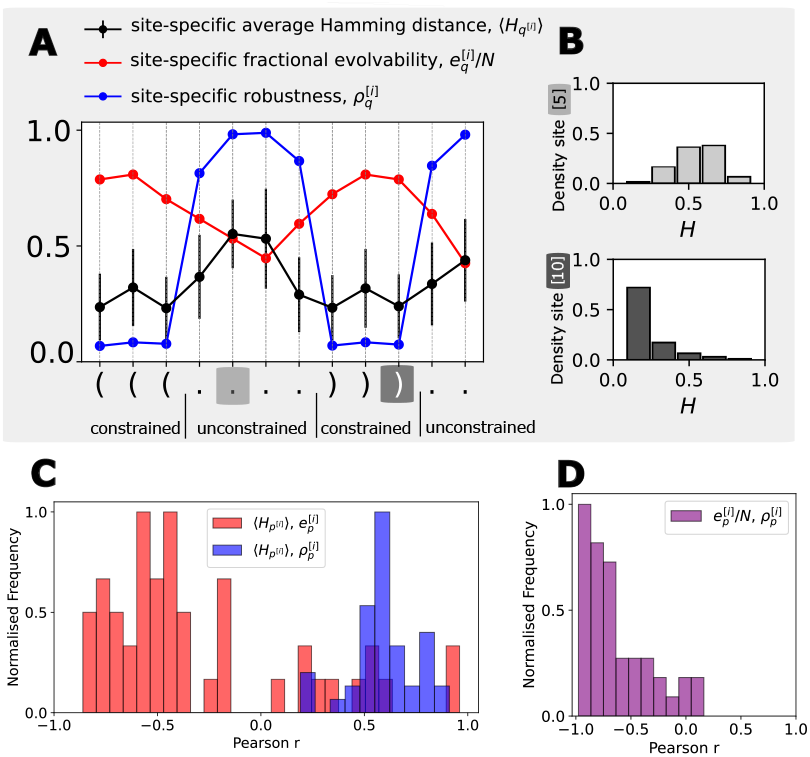
Relationships between site-specific robustness, evolvability, and phenotypic change **(A)** shows an example of the site-specific phenotypic robustness, fractional evolvability, and average Hamming distance (with no robustness or deleterious contribution), for a high frequency phenotype (shown in dot=bracket notation) in the RNA12 GP map. **(B)** Sitespecific accessible Hamming distances for unconstrained site 5 and constrained site 10 of the phenotype in (A). **(C)** Histograms of Pearson correlation coefficients comparing site-specific robustness (blue) and site-specific fractional evolvability (red) against the average Hamming distance, and against each other in **(D)**. These histograms are produced by considering all nondeleterious phenotypes in the RNA12 GP map.

If we take into account the site-specific evolvabilities shown in Fig.3A, we see that the unconstrained site 10 accesses relatively few phenotypes, but that these differ considerably from the original phenotype (see Fig.3B, top), while the constrained site 5 accesses a larger number of different phenotypes (again evidenced by the evolvability) but which are more similar to the original phenotype (Fig.3B, bottom). We show that these results hold in general by measuring the correlations between site-specific robustness, evolvability, and average Hamming distance. The Fig.3C summarises these results for the total phenotype space of RNA12 MFE GP map, and Fig.3D confirms that sitespecific quantities of phenotypic robustness and evolvability are negatively correlated, which is unsurprising as there is a clear trade-off for a single site in a similar way as the tradeoff observed for genotypic evolvability and robustness more generally, which are also negatively correlated [12].

To summarise, sequence positions that exhibit higher robustness lead to greater phenotypic change when mutated. They could therefore be described as ‘phenotypic cliffs’ in the GP map – flat in most directions, but steep in some others. Sites with lower robustness give rise to a more undulating landscape of change with a larger number of more similar alternative phenotypes.

## IV. CONCLUSION

In previous work on GP maps [20], phenotypes are usually defined as neutral (identical) or non-neutral (different) with respect to each other, but non-neutral phenotypes can be similar or very different in terms of their structure, which can have a very significant effect on fitness [21–23]. Using this as motivation to uncover a more nuanced description of GP map structure, we look at the distribution of phenotypic distances (defined in terms of dot-bracket Hamming distance) in the RNA12 GP map. First we study the structure of the GP map through the phenotypic distance framework showing that the weighted RNA12 GP map exhibits the same structural properties that are observed in a range of other GP maps [4]. Then we verify the presence of phenotypic similarity correlations and determine at the mutational probabilities with which phenotypes of different distances can be accessed in the RNA12 GP map. Both results show that similar phenotypes are closer in mutation space than expected by chance [6, 18]. Finally, we define site-specific GP map quantities and examine their relationship, which reveals that the largest phenotypic distances are accessed through the most robust sites [14].

An avenue for further research might be to explore the conditions under which natural selection might exploit the larger phenotypic changes that can arise from the mutation of robust sites. Furthermore it may also be useful to apply this framework to other biological GP maps in which phenotypic distance matters, such as the GP maps of viral proteins in the context of antigenic escape.

## V. Acknowledgements

We wish to thank Nora S Martin for useful discussions. P.G.G. acknowledges “la Caixa” Foundation (Fellowship: LCF/BQ/EU21/11890140) and King’s College, Cambridge for funding support.

## Supplementary Information

The subscript in the equations specifies whether the quantity is measured at the level of one-point mutation neighbors of a genotype *g* or of the neutral space of a phenotype *p*. The table provides formulas for both deterministic and non-deterministic cases, as established in previous studies [1].

**Figure S1.**
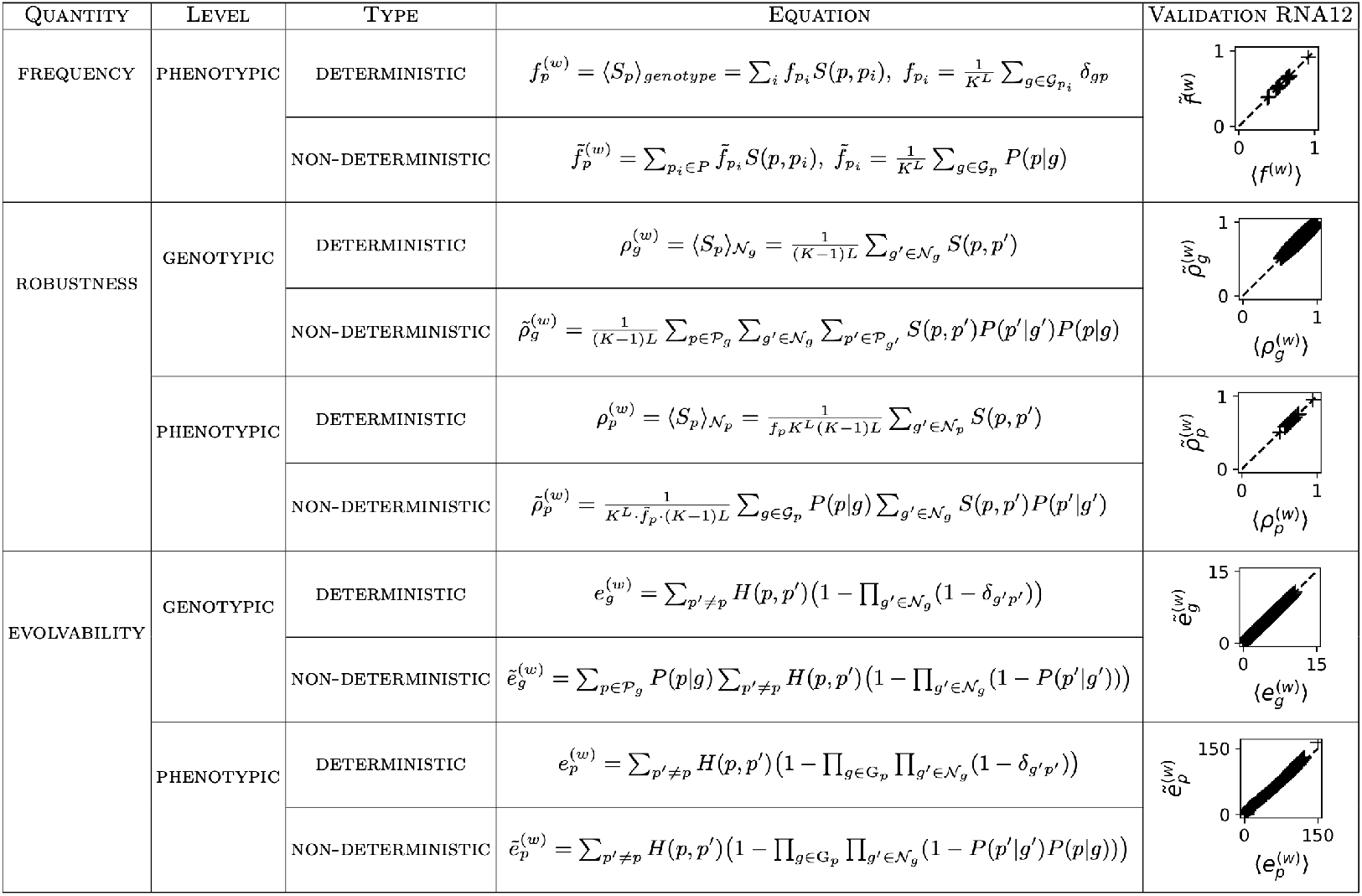
Quantities for the phenotypic distance framework in the deterministic and non-deterministic RNA12 GP map. The averages are computed with 500 samples of the RNA12 ND GP map.

The RNA ND GP map maps all possible RNA sequences of a fixed length to their respective Boltzmann ensembles of secondary structures represented in dot-bracket notation, and connects each sequence to its one-mutation neighbours to form a graph in which sequences are the nodes [1]. The phenotype associated to each genotype will be the Boltzmann ensemble found for a certain energy range (so that as Δ*E* → 0 only the MFE remains), and the Boltzmann probabilities are normalised for that set of structures [1]. Therefore, each sequence *g* folds to a secondary structure *p* with a certain Boltzmann probability *P* (*p* |*g*) = exp((*G*_*ens*_ − *G*_*p*_)*/kT*), where *G*_*ens*_ is the ensemble free energy (*G*_*ens*_ = − *kT* · ln*Z* with partition function *Z*) and *G*_*p*_ is the free energy that corresponds to structure *p* [2]. In other words, at any point in time in an RNA population of genotype *g*, the number of RNAs that fold to a certain phenotype *p* will be proportional to the Boltzmann probability *P* (*p*|*g*). In this study, we fix the RNA length to *L* = 12, so we construct the RNA12 ND GP map. For each RNA nucleotide sequence, the secondary structure ensemble, its ensemble free energy, and the free energy of each structure within the ensemble are computed using the ViennaRNA subopt [3] function set to default parameters (e.g. temperature *T* = 37^*°*^C) and energy range above the minimum free energy as Δ*E* = 15*k*_*B*_*T*, like in previous works [1].

The validation in RNA12 shows that the phenotypic framework is correctly integrated, by showing the correspondence between the non-deterministic weighted quantities in the RNA ND GP map of sequence length *L* = 12 and the averages of sampled deterministic weighted quantities [1].

## References

[1] E. Zuckerkandl and L. Pauling, Evolutionary Divergence and Convergence in Proteins, in Evolving Genes and Proteins, edited by V. Bryson and H. J. Vogel (Academic Press, 1965) pp. 97–166.

[2] H. Schwalbe, J. Buck, B. Fürtig, J. Noeske, and J. Wöhnert, Structures of RNA Switches: Insight into Molecular Recognition and Tertiary Structure, Angewandte Chemie International Edition 46, 1212 (2007).

[3] M. Doetsch, R. Schroeder, and B. Fürtig, Transient RNA–protein interactions in RNA folding, The FEBS Journal 278, 1634 (2011).

[4] S. E. Ahnert, Structural properties of genotype–phenotype maps, Journal of The Royal Society Interface 14, 20170275 (2017).

[5] Q. Vicens and J. S. Kieft, Thoughts on how to think (and talk) about RNA structure, Proceedings of the National Academy of Sciences 119, e2112677119 (2022), publisher: Proceedings of the National Academy of Sciences.

[6] L. W. Ancel and W. Fontana, Plasticity, evolvability, and modularity in RNA, Journal of Experimental Zoology 288, 242 (2000).

[7] P. García-Galindo, S. E. Ahnert, and N. S. Martin, The non-deterministic genotype–phenotype map of RNA secondary structure, Journal of The Royal Society Interface 20, 20230132 (2023).

[8] P. Schuster, W. Fontana, P. F. Stadler, and I. L. Hofacker, enFrom sequences to shapes and back: a case study in RNA secondary structures, Proceedings of the Royal Society of London. Series B: Biological Sciences 255, 279 (1994).

[9] S. F. Greenbury, S. Schaper, S. E. Ahnert, and A. A. Louis, Genetic Correlations Greatly Increase Mutational Robustness and Can Both Reduce and Enhance Evolvability, PLOS Computational Biology 12, e1004773 (2016).

[10] A. Wagner, The origins of evolutionary innovations: a theory of transformative change in living systems (OUP Oxford, 2011).

[11] A. Wagner, Robustness and Evolvability in Living Systems 10.1515/9781400849383 (2013).

[12] A. Wagner, Robustness and evolvability: a paradox resolved, Proceedings of the Royal Society B: Biological Sciences 275, 91 (2008).

[13] P. García-Galindo and S. E. Ahnert, enPhenotypic plasticity can be an evolutionary response to fluctuating environments (2024), pages: 2024.10.02.614758 Section: New Results.

[14] S. F. Greenbury and S. E. Ahnert, The organization of biological sequences into constrained and unconstrained parts determines fundamental properties of genotype–phenotype maps, Journal of The Royal Society Interface 12, 20150724 (2015), publisher: Royal Society.

[15] M. Weiß and S. E. Ahnert, enPhenotypes can be robust and evolvable if mutations have non-local effects on sequence constraints, Journal of The Royal Society Interface 15, 20170618 (2018).

[16] S. Schaper and A. A. Louis, The Arrival of the Frequent: How Bias in Genotype-Phenotype Maps Can Steer Populations to Local Optima, PLoS ONE 9, e86635 (2014).

[17] S. Wuchty, W. Fontana, I. L. Hofacker, and P. Schuster, Complete suboptimal folding of RNA and the stability of secondary structures, Biopolymers 49, 145 (1999).

[18] N. S. Martin and S. E. Ahnert, EnglishThe Boltzmann distributions of molecular structures predict likely changes through random mutations, Biophysical Journal 122, 4467 (2023), publisher: Elsevier.

[19] J. Aguirre, J.M. Buldú, M. Stich, and S. C. Manrubia, enTopological Structure of the Space of Phenotypes: The Case of RNA Neutral Networks, PLoS ONE 6, e26324 (2011).

[20] S. Manrubia, J. A. Cuesta, J. Aguirre, S. E. Ahnert, L. Altenberg, A. V. Cano, P. Catalán, R. Diaz-Uriarte, S. F. Elena, J.A. García-Martín, P. Hogeweg, B. S. Khatri, J. Krug, A. A. Louis, N. S. Martin, J. L. Payne, M. J. Tarnowski, and M. Weiß, From genotypes to organisms: State-of-the-art and perspectives of a cornerstone in evolutionary dynamics, Physics of Life Reviews 38, 55 (2021).

[21] A. W. T. Harvey, A. M. Carabelli, B. Jackson, R. K. Gupta, E. C. Thomson, E. M. Harrison, C. Ludden, R. Reeve, Na-Rambaut, S. J. Peacock, and D. L. Robertson, enSARS-CoV-2 variants, spike mutations and immune escape, Nature Reviews Microbiology 19, 409 (2021), publisher: Nature Publishing Group.

[22] N. Sanchez de Groot, A. Armaos, R. Graña-Montes, M. Alriquet, G. Calloni, R. M. Vabulas, and G. G. Tartaglia, enRNA structure drives interaction with proteins, Nature Communications 10, 3246 (2019), publisher: Nature Publishing Group.

[23] M. Mandal and R. R. Breaker, Gene regulation by riboswitches, Nature Reviews Molecular Cell Biology 5, (2004). 451

## References

1 P. García-Galindo, S. E. Ahnert, and N. S. Martin, “The non-deterministic genotype–phenotype map of RNA secondary structure”, Journal of The Royal Society Interface 20, 20230132 (2023).

2 L. W. Ancel and W. Fontana, “Plasticity, evolvability, and modularity in RNA”, Journal of Experimental Zoology 288, 242–283 (2000).

3 R. Lorenz, S. H. Bernhart, C. H. z. Siederdissen, H. Tafer, C. Flamm, P. F. Stadler, and I. L. Hofacker, “ViennaRNA Package 2.0”, Algorithms for Molecular Biology 6, 26 (2011).

